# Retinal organoids mirror CRISPR/Cas9 gene editing efficiency observed in vivo

**DOI:** 10.1101/2024.12.26.630388

**Authors:** Juliette Pulman, Hugo Malki, Paul Oudin, Ecem Aydin, Sophie Tran, Laura Visticot, Camille Robert, Anne De Cian, Marie As, Olivier Goureau, Jean-Paul Concordet, Deniz Dalkara

## Abstract

Human retinal organoids are *in vitro* 3D structures that recapitulate key molecular and structural characteristics of the *in vivo* retina. They include the presence of all essential retinal cell types including photoreceptors, making them relevant models for preclinical development of gene therapies. A critical knowledge gap exists in understanding their utility for gene editing optimization, particularly for specific genetic disorders. We assessed the potential of retinal organoids for optimizing CRISPR/Cas9-mediated gene editing, focusing on the therapeutically relevant *RHO* gene implicated in autosomal dominant Retinitis Pigmentosa (adRP). Using retinal organoids, *in vitro* HEK293T cells, and two humanized mouse models carrying different *RHO* mutations, we compared editing efficiencies. We observed that retinal organoids have lower transfection efficiency compared to HEK293T cells. Notably, they exhibited editing efficiencies more closely aligned with those found *in vivo*. We also observed similar delivery patterns of CRISPR/Cas9 tools in both retinal organoids and mouse retinas. These delivery patterns and editing efficiencies remained consistent across dual AAV systems and transiently delivered ribonucleoprotein complexes. Our findings demonstrate that retinal organoids achieve editing outcomes comparable to those observed *in vivo* underscoring their utility as part of a preclinical testing platform for genome editing, with implications for advancing gene therapy research in inherited retinal diseases.

**Graphical abstract:** 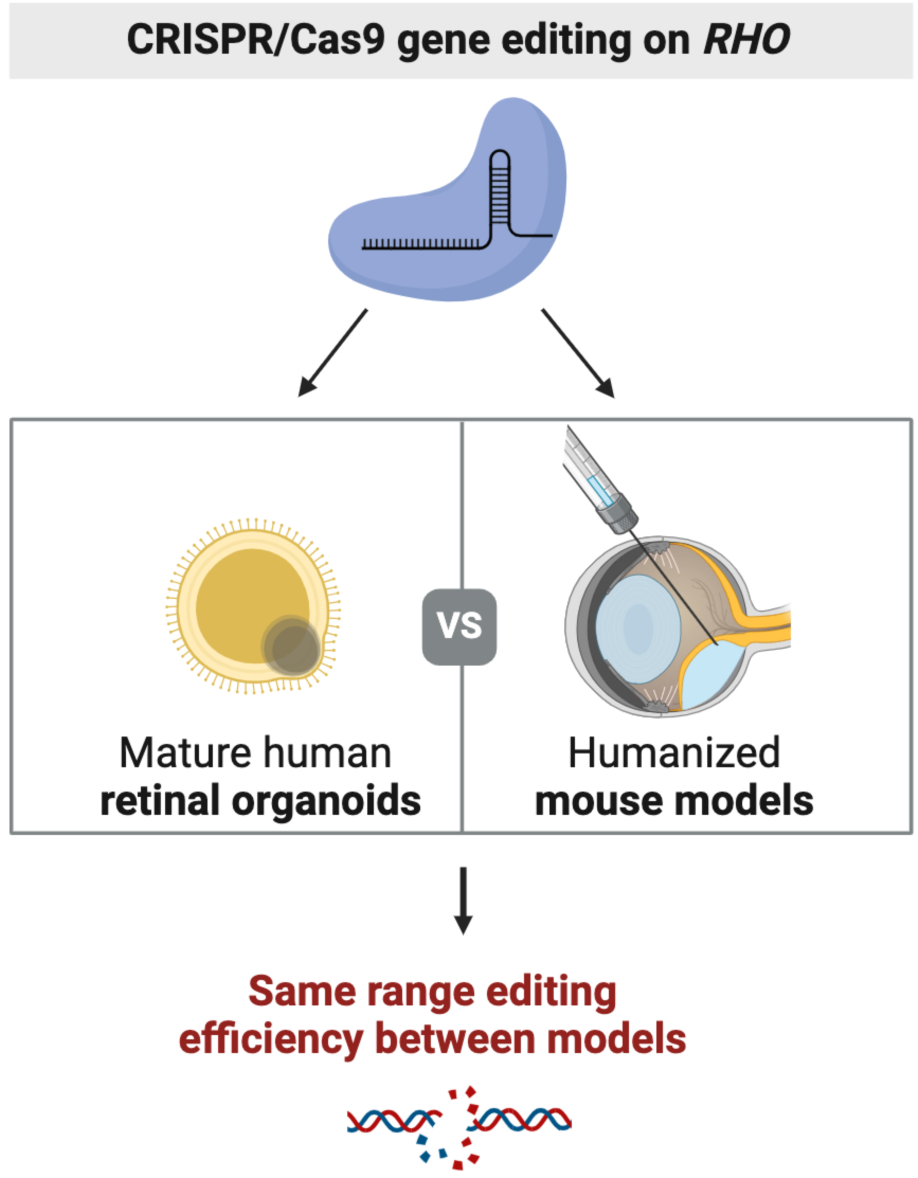

**eTOC:** Retinal organoids can be used to mirror *in vivo* mouse retina to develop CRISPR therapeutics. Here, Pulman and colleagues show similar ranges of gene editing and delivery dynamics between the organoids and *in vivo* mouse retina, highlighting the organoid’s underexplored potential for evaluating gene editing therapies in retinal diseases.

## Introduction

Organoids are lab-grown three-dimensional (3D) miniaturized structures derived from stem cells that recapitulate essential characteristics and functions of various organs. These models have become powerful tools for developmental studies, drug screening, and as a source of cells for therapeutic applications in regenerative medicine. In vision research, the differentiation of patient-derived induced pluripotent stem cells (iPSCs) into 3D retinal organoids has enabled the generation of all essential retinal cell types, including photoreceptors (Reichman et al. 2014; Zhong et al. 2014; Kuwahara et al. 2015; O’Hara-Wright and Gonzalez-Cordero 2020). These human retinal organoids have proven instrumental in modeling inherited retinal dystrophies to investigate disease and they are increasingly being leveraged to test gene therapies (Lane et al. 2020; Rodrigues et al. 2022; Boon et al. 2023; Duan et al. 2024; Kurzawa-Akanbi et al. 2024; Sladen et al. 2024). They are particularly valuable when animal models fail to recapitulate crucial disease phenotypes, such as lack of photoreceptor degeneration observed in RDH12-knock-out mice (Feathers et al., 2019) or in mouse models of type-10 Leber Congenital Amaurosis with CEP290 mutations (Maeda et al. 2006). They have gained recognition from regulatory authorities like the FDA now allowing their use in models in preclinical testing packages. By reflecting human genetics and physiology more accurately than animal models, retinal organoids serve as a critical intermediary, bridging the gap between preclinical studies and clinical trials of new drug products.

Human-relevant models are especially crucial for testing gene editing therapies, in the context of autosomal dominant retinal diseases, where gain-of-function or dominant-negative mutations demand targeted therapies. CRISPR/Cas9-based gene editing targets the specific mutations causing these conditions, and it can selectively disrupt the toxic allele through mutation-dependent or independent knockout strategies. A half dozen studies thus far have targeted the *RHO* gene implicated in adRP *in vivo*, using CRISPR/Cas9 (Giannelli et al. 2018; Li et al. 2018; Tsai et al. 2018; Wu et al. 2022 Feb 10; Yan et al. 2023; Cui et al. 2024). These preclinical advances have recently led to a phase I clinical trial for *RHO*-related RP in China (NCT05805007) conducted by Peking University Third Hospital. Yet, translating research findings into clinical success remains challenging, partly due to the lack of models that faithfully recapitulate the human photoreceptor genomic environment. Retinal organoids, with their patient-specific genetic background offer a relevant platform to screen single-guide RNAs (sgRNAs) and refine gene editing strategies for conditions like *RHO*-associated adRP. Indeed, retinal organoids not only harbor the patient’s entire genomic landscape but they can also be grow to advanced stages of differentiation where specific features such as outer segments can be observed.

Our study thus focused on using CRISPR/Cas9 to target the *RHO* gene and explore the potential of mature iPSC-derived human retinal organoids for testing gene editing in photoreceptors. Testing of gene editing tools has been performed in rodents (Wu et al. 2022 Feb 10; Qin et al. 2023; She et al. 2023; Su et al. 2023) and non-human primates (Maeder et al. 2019) but not directly in human retinal organoids. Gene editing has predominantly been applied to iPSCs prior to generation of 3D organoids in order to observe the effect of gene correction on the disease phenotype (Lane et al. 2020; Chirco et al. 2021; Rodrigues et al. 2022; Afanasyeva et al. 2023; Park et al. 2023; Siles et al. 2023). Only a single study delivered CRISPR/Cas9 to iPSC derived retinal pigment epithelium (RPE) layer using lentiviral vectors to disrupt *VEGFA* gene involved in choroidal neovascularization (Park et al. 2023). The paucity in studies using the same strategy in photoreceptors in 3D retinal organoids underscores the technical challenges in targeting photoreceptors, which harbor most causative mutations in inherited retinal dystrophies (Du et al. 2024). Overcoming the delivery barrier thus remains a central obstacle to effective gene editing in human photoreceptors, especially when it comes to the delivery of gene editing tools. Indeed, mRNA encoding GFP has been successfully delivered into photoreceptors of retinal organoids using lipid nanoparticles (LNP), generating detectable levels of fluorescent protein but the same strategy showed limited efficiency, for CRISPR/Cas9 delivery (Eygeris et al. 2024 May 13). We recently achieved therapeutically relevant gene editing efficiencies *in vivo* by direct delivery of CRISPR/Cas9 ribonucleoprotein (RNP) into mouse retinas. Our in vivo gene efficiency with direct RNP delivery was comparable to the editing efficiencies obtained using Adeno-Associated Virus (AAV) mediated CRISPR/Cas9 delivery (Pulman et al., 2023). These advancements motivated us to compare these delivery strategies in 3D retinal organoids to understand their value in predicting gene editing outcomes.

In this study, we compared for the first time the efficiency of CRISPR/Cas9-mediated gene editing in photoreceptors of mature human retinal organoids and mouse retinas to determine whether organoids can reliably predict *in vivo* outcomes. Our findings demonstrate that human retinal organoids, containing mature photoreceptors embedded in a 3D environment closely mimick the *in vivo* retina, allowing us to achieve gene editing efficiencies close to those observed *in vivo*. This work establishes a crucial foundation for using organoids to optimize CRISPR/Cas9 delivery strategies, ultimately contributing to the development of more effective genome editing therapies for human inherited retinal dystrophies.

## RESULTS

### 1/ An efficient *RHO*-targeting sgRNA is identified through *in vitro* screening

To edit the *RHO* gene using CRISPR/Cas9 for potential future use in gene therapy, we designed sgRNAs targeting the human *RHO* sequence across the first three of five exons. The choice of position along the coding sequence followed the rules of the mRNA-mediated decay system, which identifies premature termination codons in transcripts, to ensure efficient gene knockout relevant for therapy (Kurosaki et al. 2019). sgRNAs were ranked by the Doench’16 efficiency score (Doench et al. 2016) using the CRISPOR tool (Concordet and Haeussler 2018), then the best guides were selected for each exon. We selected nine sgRNAs: five targeting *RHO* exon 1, three targeting *RHO* exon 2, and one targeting *RHO* exon 3 (Fig 1A). We first performed an *in vitro* screening in the human HEK293T cell line. Each sgRNA was synthesized *in vitro,* complexed individually with the SpCas9-GFP enzyme and then transfected into HEK293T cells. For comparison, we included a sgRNA targeting *VEGFA* that has already been shown to be efficient both *in vitro* and *in vivo* in mouse retinas.

**Figure 1.**
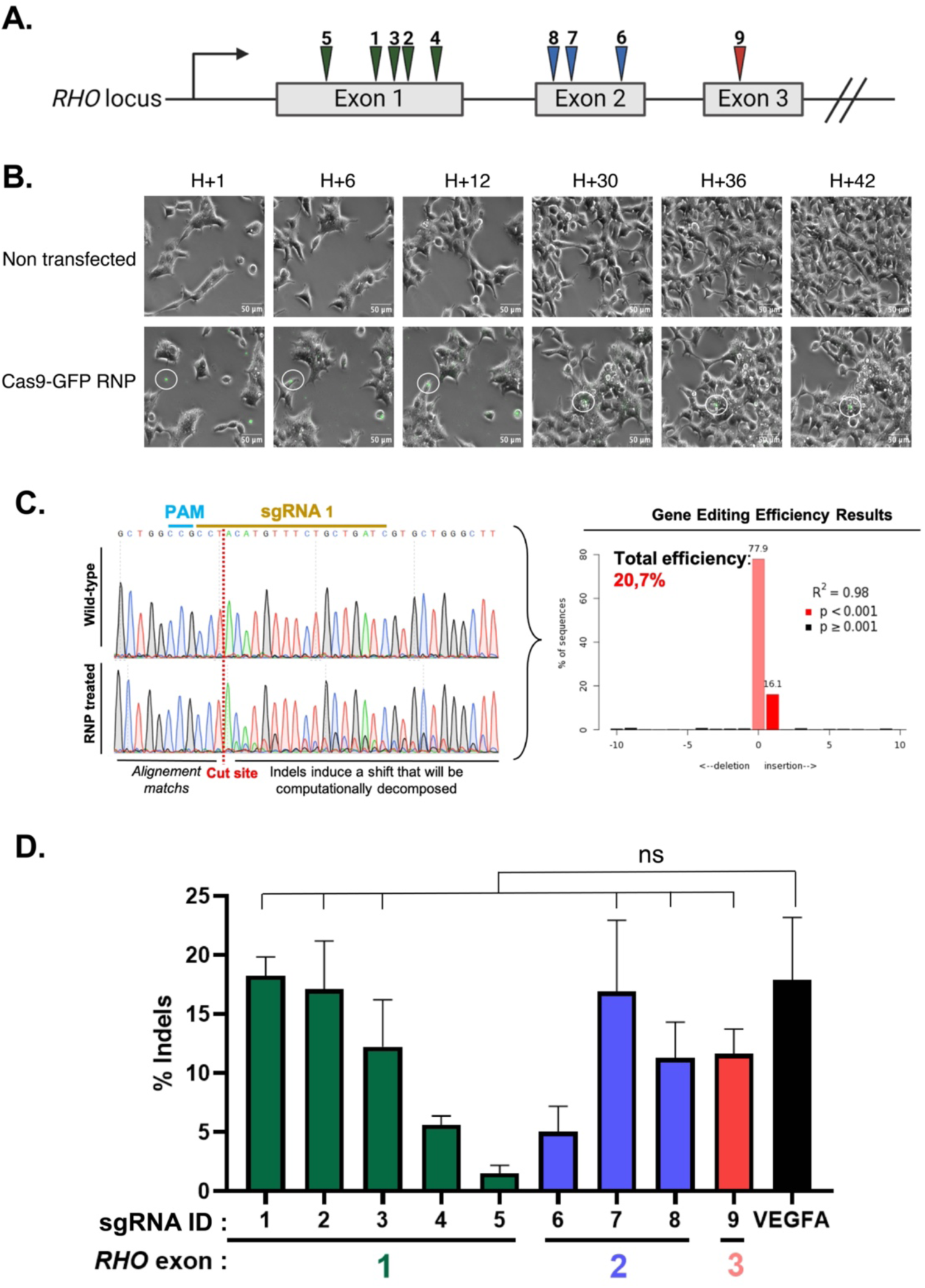
Efficient sgRNA for targeting *RHO* identified through *in vitro* CRISPR/Cas9 transfection. **A.** The different sgRNA tested, positioned on the *RHO* locus across the 3 first exons**. B.** Timeline of a HEK293T cells transfection of Cas9-RNP/Lipofectamine complexes (in green). An aggregate of RNP was circled to follow its path. **C.** TIDE analysis of sequenced amplicon, example of sgRNA 1. The algorithm compares untreated mock control with RNP treated cells chromatograms to determine a percentage of edition (eff – in TIDE OUTPUT) by decomposing the file of treated condition. **D.** Result of *in vitro* screening of the 9 sgRNA tested on HEK293T compared to a previously published sgRNA targeting *VEGFA* (Kim et al. 2017) Mean ± SEM. Ordinary One-way ANOVA test, Dunnett’s multiple comparisons test showing that 6 sgRNAs have similar editing efficiency that sgRNA *VEGFA*. ns= not significant.

To observe the dynamics of RNP transfection and cellular interactions, we captured images of the lipofection process over 42 hours. Throughout the experiment, we observed that Cas9-GFP RNP/Lipofectamine 2000 complexes gradually sedimented onto the surface of the plate. As the HEK293T cells grew and divided, they moved toward these RNP/Lipofectamine complexes. By the end of the experiment, only a few remaining GFP-positive complexes were visible, suggesting that most of the RNPs were internalized and degraded by the cells. This dynamic movement is characteristic of cells growing in 2D (Fig 1B and Sup Video S1 and S2) and likely introduces a bias in the way gene editing tools access the cell’s nuclei compared to in vivo conditions.

Next, to evaluate the editing efficacy of our lipofection, we used the TIDE assay to determine the efficacy of each sgRNA. The TIDE assay determines the editing frequency generated in a pool of cells using Sanger sequencing (Brinkman et al., 2014). As shown in the example of sgRNA 1, we observed the generation of mutations around the cut site of our sgRNA compared to control non-transfected HEK293T cells (Fig 1C). We analyzed all our tested sgRNAs and identified six sgRNAs targeting *RHO* as efficiently as the *VEGFA* sgRNA (Fig 1D). Among these, sgRNA 7 was randomly selected for further experiments on retinal organoids.

### 2/ Cas9 RNP delivery into human retinal organoids achieves RHO editing rates comparable to in vivo delivery into a human RHO knock-in mice

Retinal organoids derived from human iPSCs spontaneously self-organize in 3D and develop structures naturally present in the eye’s retina, including specialized retinal neurons such as the photoreceptors (Fig 2A). Photoreceptors consist of a nucleus, an inner segment, and an outer segment. The outer segment captures light through the concerted action of the proteins involved in phototransduction. For all our experiments, we used 150-day-old organoids that already contain mature photoreceptors and outer segment-like structures harboring the phototransduction proteins. Bright-field imaging confirmed the intact structure and lamination of the retinal organoids, as well as the presence of cilia (inner and partial outer segment) on the surface, indicative of photoreceptor differentiation (Fig 2B).

**Figure 2.**
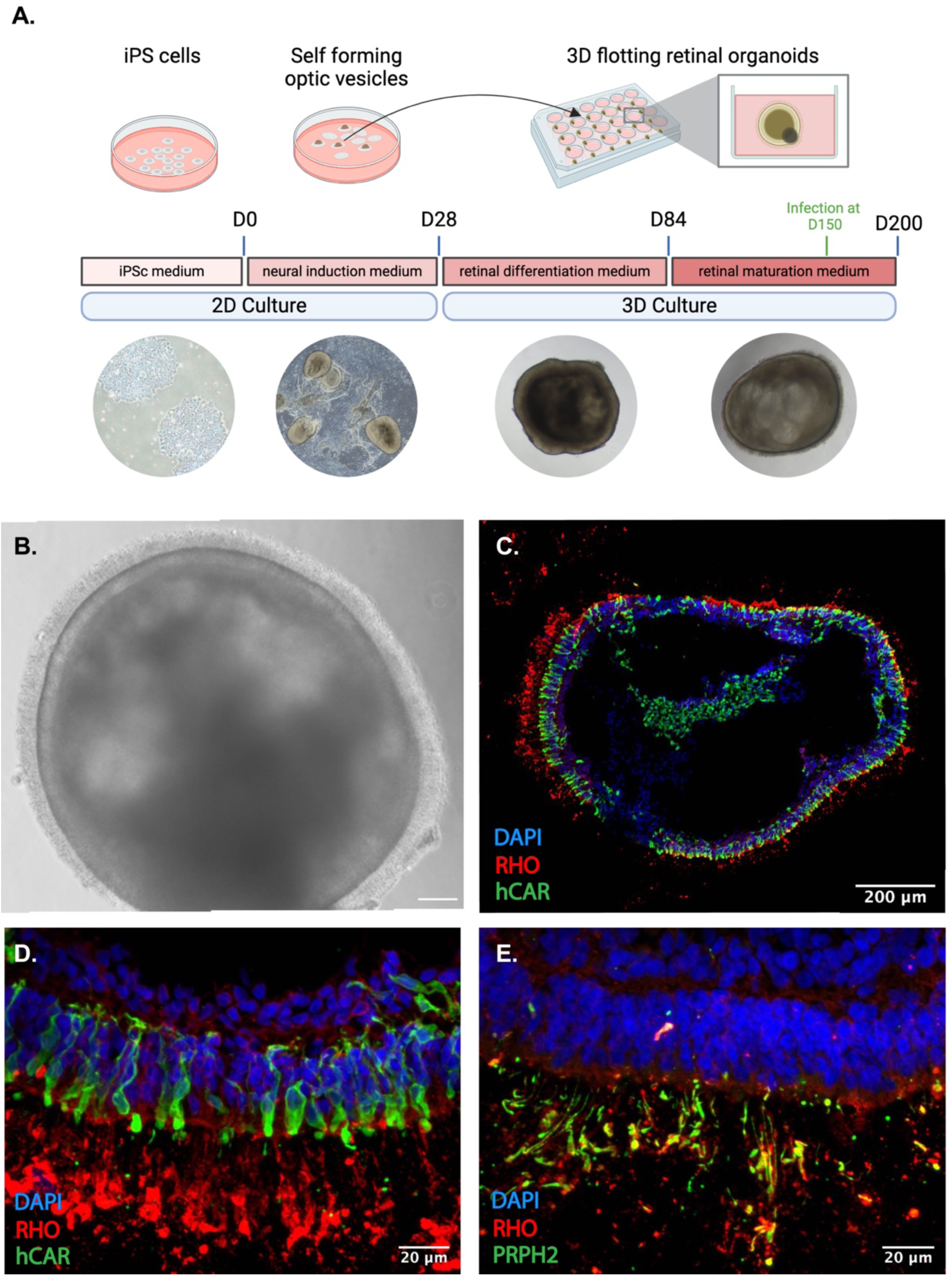
Retinal organoids exhibit lamination and photoreceptor segment development. **A.** Schematic timeline of the differentiation process going from iPSCs to mature retinal organoids. **B.** Bright-field image of a control retinal organoid at differentiation day (DD) 150 in culture showing the outer nuclear layer (ONL) and the presence of cilia on the surface, indicative of photoreceptor differentiation. Scale bar: 100 µm. **C.** Cryosection of an organoid stained with rhodopsin (*RHO*), a rod marker, and human cone arrestin (*hCAR*), a cone marker. **D.** High-magnification view of a region from (C), showing a photoreceptor arrangement highly reminiscent of the native retinal architecture. **E.** Cryosection of an organoid stained with rhodopsin and peripherin (*PRPH2*), a marker of outer segments.

Immunostaining on cryosections of an organoid stained with rhodopsin (*RHO*), a rod cell marker, and human cone arrestin (*hCAR*), a cone cell marker, illustrates the proper lamination with both rod and cone photoreceptors forming the outermost layer (Fig 2C and D). Also, immunostaining of an outer segment marker, Peripherin (*PRPH2*), showed a co-localization with rhodopsin and confirmed the presence of photoreceptor outer segment-like structures at the organoid surface (Fig 2E). The presence of both rods and cones containing outer segment-like structures allows us to test CRISPR/Cas9 delivery in a 3D context that closely mimics that of *in vivo* mature photoreceptors.

Since retinal organoids are an *ex vivo* model, we initially applied the CRISPR/Cas9 transfection conditions used for HEK293T cells directly to the organoids. We adjusted the concentration of Cas9 RNP complexed with Lipofectamine 2000 to 60 nM to maintain a consistent ratio of Cas9 RNP per cell across experiments. However, at this concentration, we could not detect any indels in the retinal organoids compared to the non-transfected ones (Fig 3A), indicating that these complex structures have a lower transfection efficiency than 2D dividing HEK293T cells.

**Figure 3.**
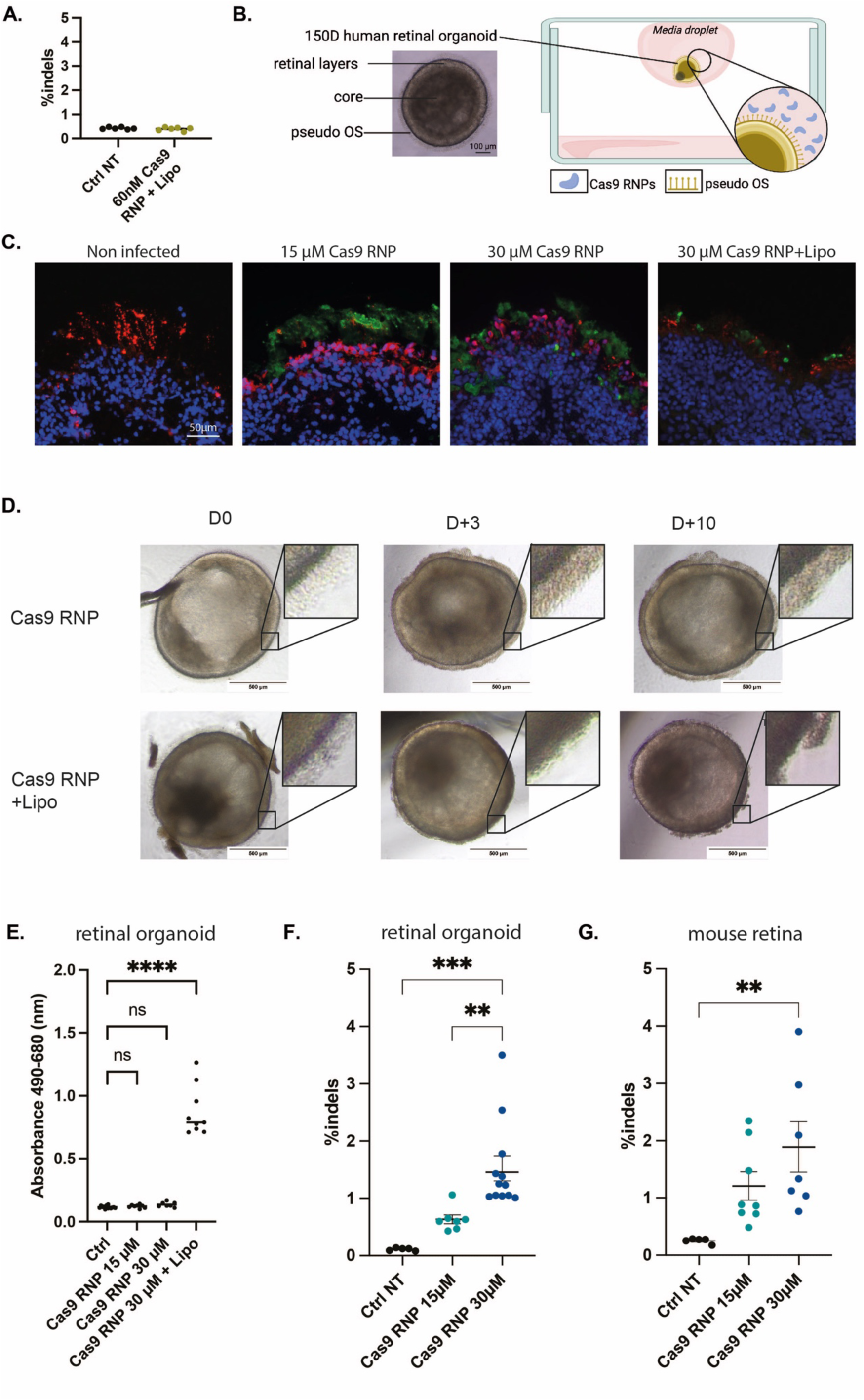
CRISPR/Cas9 RNP delivery to retinal organoids displays similar editing efficacy to *in vivo* subretinal injections. **A.** Indels in retinal organoids after transfection of Cas9-RNP/Lipofectamine complexes at the same ratio used for HEK293T transfections. NGS analysis was performed 7 days after transfection. Each dot represents one organoid. **B.** Schematic representation of the hanging drop strategy to deliver Cas9 RNP to the retinal organoids. **C.** Human retinal organoids cross-sections staining of nuclei (DAPI, blue), photoreceptors (Recoverin, red) and Cas9 protein (green). Organoids were collected three days after treatment of 15 or 30 µM Cas9 RNPs or 30 µM Cas9-RNP/Lipofectamine. **D.** Bright-field image of retinal organoids treated with Cas9 RNP or Cas9-RNP/Lipofectamine at DD150 (D0), 3 or 10 days after (D+3 or D+10) **E.** Cell cytotoxicity analysis through LDH measurement of human retinal organoids after Cas9 RNP +/- Lipofectamine 2000 treatment. Mean ± SEM. Ordinary One-way ANOVA test, Dunnett’s multiple comparisons. ****: pvalue<0.0001. ns: non-significant **F.** Indels in retinal organoids after transfection of 15 or 30 µM CRISPR/Cas9 RNP. NGS analysis was performed 7 days after transfection. Each dot represents one organoid. Mean ± SEM. Ordinary One-way ANOVA test, Dunnett’s multiple comparisons. **: pvalue<0,005. ***: pvalue<0.0005. **G.** Indels in h*RHO*.p347S mouse retina after 1 µL subretinal injection of15 or 30 µM CRISPR/Cas9 RNP. NGS analysis was performed 7 days post-injection. Each dot represents the whole neural retina isolated from a single mouse eye. Mean ± SEM. Ordinary One-way ANOVA test, Dunnett’s multiple comparisons. **: pvalue<0,005.

To investigate if retinal organoids mimic *in vivo* conditions, we also tested Cas9 RNP delivery without any vector as described in our previous studies (Pulman et al. 2024). We first confirmed that the culture medium of the organoids was not affecting the proper formation of the Cas9 RNP. Dynamic light scattering showed a homogeneous and unchanged hydrodynamic size of the Cas9 RNP in the culture medium (Fig S1). When adding Lipofectamine 2000, however, we observe significant aggregations (Fig S1). As a 30 μM concentration is challenging to achieve in a 100 μL of culture medium volume, we implemented a hanging drop (HD) strategy, allowing to transfect retinal organoids using only 6 µL of media (Fig 3B).

Three days after exposing the retinal organoids to Cas9 RNPs wither alone or complexed with Lipofectamine, we observed that the Cas9 proteins were blocked in the outer segment-like structure of the photoreceptors, in the periphery of the organoids for both 15 and 30 μM Cas9 RNP (Fig 3C). We also observed a degradation of the outer segment-like structure in the organoids treated with the Cas9 RNP/Lipofectamine complexes, with less internalized Cas9 protein (Fig 3C). We confirmed Lipofectamine related toxicity using bright-field images, which showed disorganization of the outer segment-like structures in organoids treated with Cas9 RNP/Lipofectamine complexes. (Fig 3D). We also performed a cell cytotoxicity assay based on the measurement of the lactate dehydrogenase (LDH) in the culture medium revealing no cytotoxicity with Cas9 RNP alone while revealing significant toxicity in those conditions including Lipofectamine 2000 (Fig 3E). We therefore noted a similar response to Lipofectamine mediated toxicity in vivo and in organoids and discontinued experiments with Lipofectamine 2000.

We concentrated our efforts on investigating editing efficiency of the Cas9 RNP complexes coupled to our *in vitro* selected sgRNA 7. We observed a dose-dependent editing on the *RHO* gene with these RNPs achieving a 1.5 % indel efficiency when treated with 30 μM of naked Cas9 RNP (Fig 3F). Finally, we compared this efficacy to *in vivo* editing using the same conditions in a humanized mouse model with the p.P347S mutation integrated at a heterozygous stage (Figure S2). In this model, the murine Rho remains present and unchanged (Li et al. 1996). In vivo, in the h*RHO*.P347S mouse model, we observed a similar editing efficiency of 2% indels (Fig 3G).

### 3/ Dual-AAV mediated CRISPR delivery to retinal organoids mirrors *in vivo* gene editing outcomes in two distinct mouse models of adRP

To further evaluate the response of retinal organoids to gene editing tools compared to *in vivo* mouse retina, we tested the most common retinal delivery method of dual AAVs. Wu et al. described an AAV construct with the sCMV promoter driving expression of SpCas9 and proved its efficacy in editing photoreceptor cells in mouse retinas in vivo (Wu et al., 2022). We used the same dual AAV strategy, with one AAV containing the SpCas9 and the other containing the sgRNA and a GFP reporter expression cassette (Fig 4A). As SpCas9 cDNA is relatively big (4.1 kb), the expression cassette with sCMV slightly exceeds the AAV packaging capacity (usually ∼4.7kb). Therefore, we first quality checked the SpCas9 AAV8 production using dynamic light scattering showing 91% full capsids and a homogenous hydrodynamic size of 29.4nm (Fig. S3).

**Figure 4.**
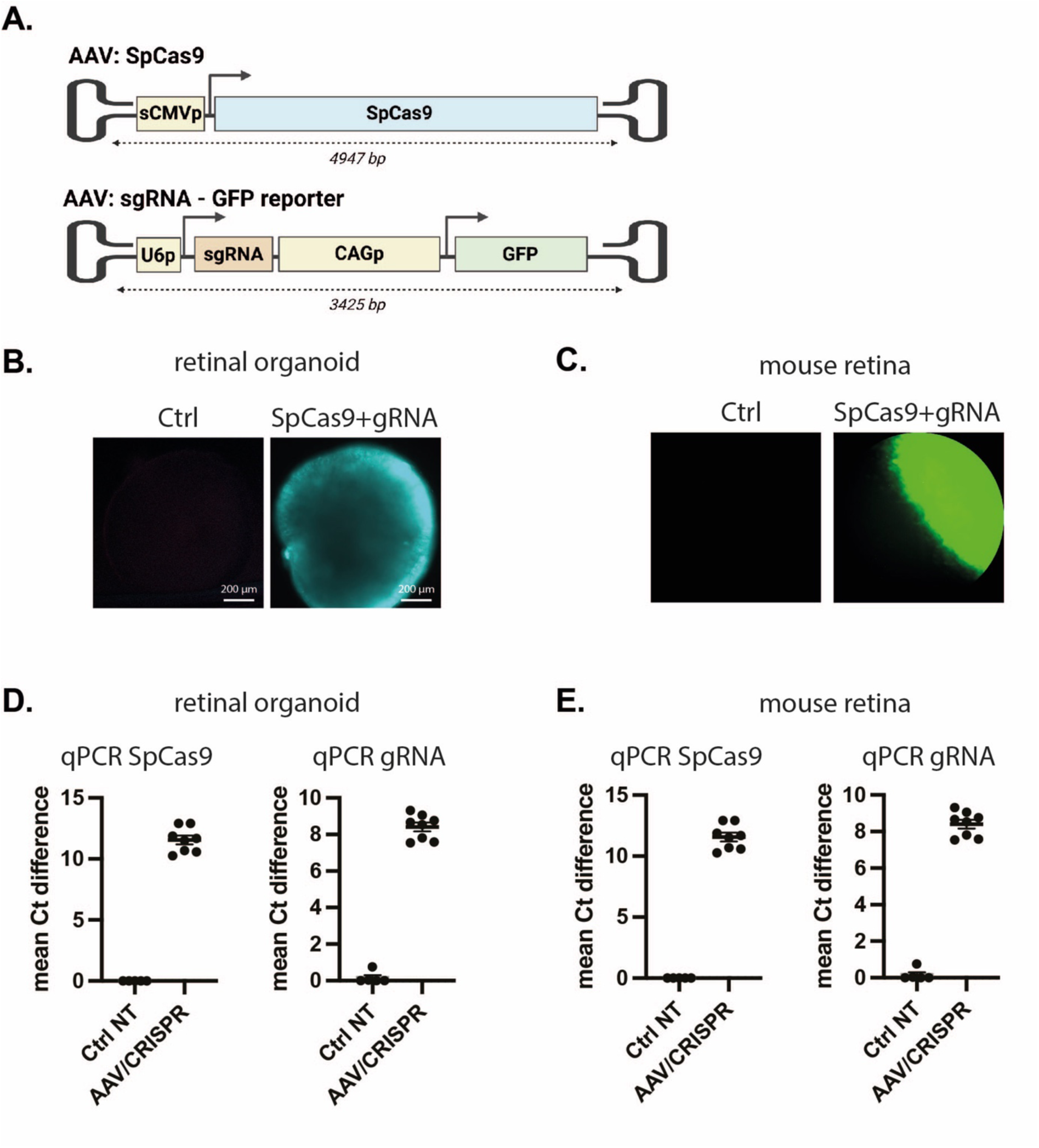
Dual-AAV CRISPR/Cas9 is expressed in human retinal organoids and *in vivo*. **A.** Double AAV strategy used to deliver the CRISPR/Cas9 system, inspired from Wu et al. (Wu et al. 2022 Feb 10) One AAV vector carries a SpCas9 cDNA driven by the sCMV promoter. One AAV vector carrying the h*RHO*-specific sgRNA (sgRNA 1, 7, or from Wu and colleagues)-expressing cassettes and a GFP cDNA driven by CAG promoter. This illustration does not reflect actual scale. **B.** GFP expression in retinal organoids 7 days after transduction with dual-AAV CRISPR. Scale bar: 200 μm. **C.** GFP expression in fundus imaging of h*RHO*.P347S mouse eye 14 days after subretinal injection. **D&E.** Semi-quantitative expression level of exogenous SpCas9 or sgRNA after transduction with dual-AAV CRISPR at a total dose of 5E9 vg and untransduced controls. **D.** in retinal organoids. **E.** in h*RHO*.P347S mouse retinas.

We then transduced in parallel retinal organoids and mouse retinas, using the dual-AAV CRISPR strategy. We injected the h*RHO*.P347S mice and a second *hRHO* mouse model carrying the p.P23H mutation subretinally with the dual AAVs. The p.P23H model harbors one humanized mutated allele replacing one wild-type murine allele and is fused to the fluorescent protein RFP (Robichaux et al. 2022). We confirmed GFP expression, from the AAV carrying the sgRNA, in retinal organoids and the h*RHO*.P347S mouse retina by immunofluorescence (Fig. 4B and C). We also confirmed SpCas9 and the sgRNA expression via semi-quantitative qPCR analysis (Fig. 4D and E). Our results indicate successful expression of SpCas9 and sgRNA in both models.

We analyzed the distribution of AAV after infections by assessing GFP expression on cryosections of retinal organoids and mouse retinas. In retinal organoids, a strong GFP expression was observed in the outermost layer, corresponding to the localization of recoverin-positive photoreceptors (Fig. 5A). In both mouse models, GFP expression was restricted to the photoreceptor layer (Fig. 5B and 5C). We observed significant differences in the rate of retinal degeneration between the two mouse models, a much faster outer nuclear layer (ONL) loss seen in the h*RHO*.P347S model compared to the h*RHO*.P23H model (Fig. 5B and 5C).

**Figure 5.**
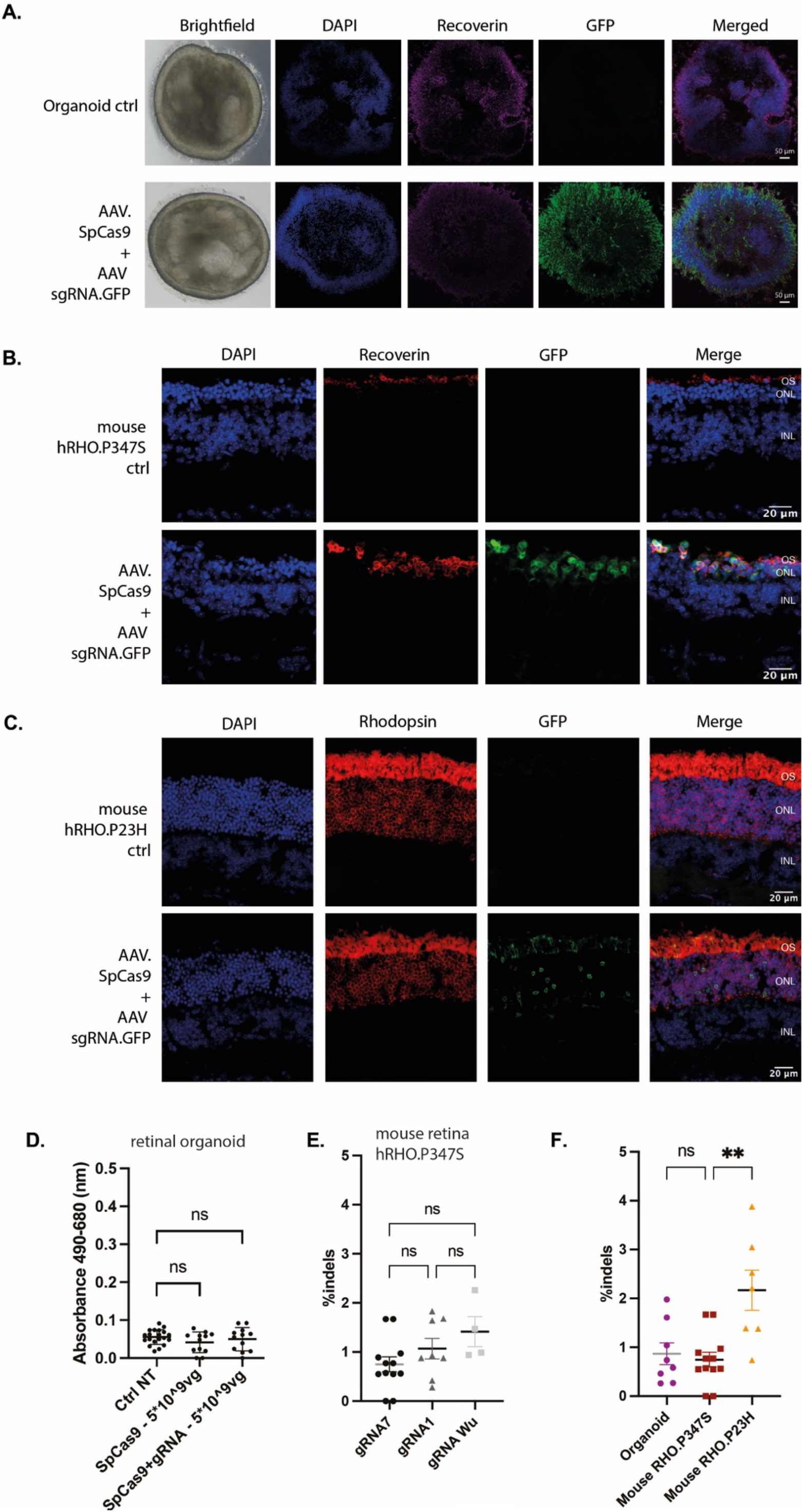
CRISPR/Cas9 AAV delivery to retinal organoids displays similar editing efficacy to *in vivo* subretinal injections. **A.** Cryosection of an organoid 21 days after dual-AAV CRISPR transduction, stained with Recoverin, a photoreceptor marker. **B.** Cryosection of neural retina sections of mouse h*RHO*.P347S stained for nuclei (DAPI, blue), photoreceptors (Recoverin, Red) and GFP (green). **C.** Cryosections of neural retina of mouse h*RHO*.P23H stained for nuclei (DAPI, blue), GFP (green) and photoreceptors expressing human rhodopsin (RFP, red). OS: outer segment, ONL: outer nuclear layer, INL: inner nuclear layer. **D.** Cell cytotoxicity analysis through LDH measurement of retinal organoids after dual-AAV CRISPR transduction using only the AAV carrying SpCas9 or the dual-AAV CRISPR at a total dose of 5E9 vg. ns: non-significant. **E.** Indels in h*RHO*.p347S mouse retina following dual-AAV CRISPR subretinal injection with different sgRNAs. sgRNA1 and sgRNA7 were identified in our in-house screen; sgRNA “Wu” is designed from the two publications from Tsai, Wu and colleagues. (Tsai et al. 2018; Wu et al. 2022 Feb 10) Each dot represents the whole neural retina isolated from a single mouse eye. Mean ± SEM. Ordinary One-way ANOVA test, Dunnett’s multiple comparisons. ns: non-significant. **F.** Indels in retinal organoids, h*RHO*.p347S mouse retina or h*RHO*.p23H mouse retina after transduction of dual-AAV CRISPR using sgRNA 7. Each dot represents one organoid or the whole neural retina isolated from a single mouse eye. Mean ± SEM. ns: non-significant. **: pvalue<0,005.

We assessed the cytotoxicity of the dual AAV strategy in retinal organoids. At a dose of 5E9 vg per organoid, no significant increase of LDH levels in culture media was observed (Fig. 5D), indicating minimal cytotoxicity. To validate the efficiency of our selected sgRNA 7, we compared its performance to a previously reported gRNA from Wu and colleagues. (Tsai et al. 2018; Wu et al. 2022 Feb 10). Additionally, we included a second sgRNA from our initial panel for comparison (sgRNA 1). Each of the three sgRNAs was injected separately into h*RHO*.P347S mouse retinas, and the editing efficiency was assessed. No significant difference in editing efficiency was observed among the three sgRNAs (Fig. 5E). Based on these findings, we proceeded with sgRNA 7 for subsequent experiments.

Finally, we compared the editing efficiency of the dual-AAV CRISPR approach in retinal organoids and mouse retinas. Editing efficiencies were found to be in the same range between the retinal organoids and the mouse retinas (Fig. 5F). It is important to note that the editing efficiency varied between h*RHO*.P23H and h*RHO*.P347S mouse retinas (Fig. 5F), likely due to the faster degeneration rate observed in the h*RHO*.P347S model allowing for editing in fewer surviving photoreceptors. Altogether, our results demonstrate dual-AAV CRISPR editing efficacy in mature retinal organoids, just like editing rates using RNP occurs at a comparable rate to those observed in mouse retinas harboring the *hRHO* gene. The gene editing rates remain similar between the human retinal organoids and mouse models with mutant *hRHO* knock in albeit differences in the number of surviving photoreceptors skewing the results for the mouse line with the faster degeneration rate.

## Discussion

In this study, we investigated whether 3D human retinal organoids can serve as a predictive model for testing gene editing efficacy, complementing the mouse models. While *in vivo* models are valuable, human gene sequences can only be tested after integration in the animal model’s genome and replication of human-specific chromatin accessibility, epigenetic states, and transcriptional landscapes of the human sequence in animal model photoreceptors remain limited. Indeed, those parameters are important for evaluating the efficiency of sgRNAs. Human immortalized cell lines harbor the human genome but have different gene expression and chromatin context and cannot be used to model human photoreceptors. (Chakrabarti et al. 2019; Moreb and Lynch 2021) A complementary model, such as retinal organoids, provides a platform that closely mimics human genetic context, allowing researchers to assess sgRNAs targeting efficiency with greater relevance to clinical scenarios. Additionally, such a model enables the testing of off-target effects or genomic translocations potentially induced by Cas9 within a human genome, which cannot be evaluated in rodents. By bridging the gap between *in vitro* and *in vivo* approaches, complementary models like retinal organoids greatly facilitate the optimization of gene editing strategies, enhancing the likelihood of successful therapeutic development for inherited retinal dystrophies.

Retinal organoids were primarily used for developmental studies or drug screening and are being increasingly used for gene therapy development. (Lane et al. 2020; Boon et al. 2023; Duan et al. 2024; Kurzawa-Akanbi et al. 2024) Also, anti-sense oligonucleotides have been directly tested on mature organoids. (Dulla et al. 2018; Kaltak et al. 2023) Surprisingly, they have been under-explored for testing CRISPR therapeutics. Our study demonstrates that retinal organoids can help fill this gap, particularly for assessing not only gene editing efficiency but also the delivery and safety of CRISPR tools.

Our study demonstrates that similar editing rates are achieved in human retinal organoids and mouse retinas. We treated directly the retinal organoids with naked Cas9 RNP or AAVs encoding Cas9 and compared it to mouse retina subretinal injections. The editing efficiency in retinal organoids was much closer to the in vivo results than the results in HEK293T cells. Also, we observed that delivery in 2D HEK293T cells can benefit from the sedimentation of the RNP complexes, which will not be the case for 3D floating retinal organoids. Moreover, we see a difference in the editing efficacy between our two humanized mouse models, suggesting that evaluating the editing in humanized mouse models might give misleading results compared to future human use as the mouse models degenerate at a much quicker rate shrinking the number of edited photoreceptors.

Beyond the mutation type, other factors such as the chromatin accessibility at the integration site of the human transgene and its epigenetic landscape could impact editing efficiency. For example, regions of heterochromatin or densely packed chromatin might limit the accessibility of Cas9 RNP, thereby reducing its ability to induce double-strand breaks. (Daer et al. 2017; Uusi-Mäkelä et al. 2018; Pierce et al. 2021) Additionally, the localization of the transgene within the genome could subject it to positional effects, such as differential transcriptional activity or interaction with regulatory elements, further influencing the outcome of degeneration and/or gene editing. The number of integrated transgene copies might also introduce variability, as higher copy numbers could alter the stoichiometry of editing events. Together, these factors underscore the complexity of using poorly characterized mouse models to predict editing efficiency and highlight the necessity of complementary models for a more precise evaluation.

In addition to testing the editing efficiency, retinal organoids provide a valuable platform for testing the delivery dynamics with different vectors. We showed in a previous work that subretinal injection of naked Cas9 RNP led to RNP being predominantly blocked at the level of photoreceptors’ outer segments which may represent a physical barrier during non-viral delivery of tools like Cas9 RNP (Pulman et al. 2024). Here, we found a similar pattern in retinal organoids treated with Cas9 RNP. Such findings emphasize the need for complementary models like retinal organoids to explore these physical barriers under controlled experimental conditions for therapy development. Along the same line, we conclude that the retinal organoids also represent an interesting model to evaluate cytotoxicity. For example, Dorgau and colleagues used retinal organoids to quantify the toxicities of already licensed drugs with known responses using single-cell RNA sequencing and immunofluorescence assays. (Dorgau et al. 2022) Also, the toxicity of different products including pesticides, flame retardants and other typical environmental pollutants has been assessed in organoids, confirming their potential as a retina model (Kurzawa-Akanbi et al. 2024). In our case, our previous work demonstrated the inherent toxicity of Cas9 RNP mixed with Lipofectamine 2000 as non-viral vector in wild-type mouse leading to photoreceptors outer segments damage and shedding. (Pulman et al. 2024) Here, we observed a similar cytotoxicity in retinal organoids. We observed absence of the OS-like structures, which were likely damaged and lost due to Lipofectamine reducing the Cas9 RNP signal in immunostaining.

However, it is important to note that retinal organoids still lack key features of the *in vivo* retina, such as vascularization and an immune system. Emerging technologies are rising to address these limitations. Advances like organoids-on-a-chip and co-cultures with RPE or glial cells offer promising solutions to create more physiologically relevant models. (Achberger et al. 2019; Achberger et al. 2021) These improvements could further enhance the utility of organoids demonstrated in our study, by enabling more accurate evaluation of gene editing tools in a microenvironment that better mimics human retinal physiology, including intercellular interactions and immune responses. The parallel between the editing efficiency should be taken with caution as the number of targeted cells, the AAV serotype, and the surrounding environment were different between retinal organoids and mouse retina. However, it is interesting to note the similarity in the range of editing, suggesting that it could be a good model to test CRISPR editing efficiency, to screen human sgRNA efficacy and genotoxicity on a human genetic background and in primary photoreceptors neurons, or different vectors efficiency and cytotoxicity.

Our study opens the way for using mature retinal organoids as a model to develop new gene editing therapies. The next step will be to investigate the potential of treatment of gene editing tools on a degenerative model of retinal organoids from patients with a specific mutations. We believe human retinal organoids are a robust and versatile model for testing CRISPR/Cas9-based gene editing and they will allow for comprehensive evaluation of editing efficiency, delivery dynamics, and cytotoxicity in a human-specific genetic background. By bridging the gap between traditional *in vitro* systems and *in vivo* models, retinal organoids hold significant potential to accelerate the development and optimization of gene editing therapies for inherited retinal diseases

## Materials and Methods

### sgRNA design

sgRNAs targeting human *RHO* gene were designed using CRISPOR (http://crispor.gi.ucsc.edu/). The sgRNA called “Wu” was designed according to the two publications of Tsai, Wu and colleagues (Tsai et al. 2018; Wu et al. 2022 Feb 10). The sgRNA targeting the vascular endothelial growth factor A (Vegfa) gene was published by Kim and colleagues. (Kim et al. 2017) All sgRNAs were synthesized and purified using the GeneArtTM Precision gRNA Synthesis Kit (Invitrogen), according to the manufacturer’s protocol. sgRNAs eluted in water were aliquoted and stored at -80°C. All 20 bp sequences targeted by each sgRNA are listed in Table S1.

### SpCas9 nuclease

For experiments on HEK293T cells, Streptococcus pyogenes Cas9 (SpCas9) nuclease (Aldevron) was used. For HEK293T cells live imaging lipofection, SpCas9 nuclease fused with GFP was used. For experiments on organoids and *in vivo*, SpCas9 nuclease with 2 nuclear localization sequences (NLS) (one on its N and one on its C terminal) was produced as previously described (Ménoret et al. 2015)and kept at -80°C until use.

### Cell culture

Briefly, human embryonic kidney HEK293T cells were cultivated in Dulbecco’s Modified Eagle’s Medium (DMEM, Gibco) complemented with fetal bovine serum (Gibco) at 10% and 1% penicillin-streptomycin antibiotics (Gibco). Cells were incubated at 37°C, 5% CO2.

The hiPSC-5F were cultured as our previously established protocol. (Slembrouck-Brec et al. 2019) Briefly, iPSC colonies were kept in culture in an incubator (RWD D180 Incubator) at 37 °C, under 5% CO2/95% air atmosphere, 20% oxygen tension and 80%–85% humidity. Colonies were cultured with antibiotic free mTeSRTM1 medium (StemCell Technologies), in culture dishes coated with truncated recombinant human vitronectin (StemCell Technologies) and were passaged once a week

### Retinal organoid differentiation

Organoid generation was based on our previously established adherent hiPSC differentiation. (Reichman et al. 2017; Slembrouck-Brec et al. 2019) hiPSCs were expanded to confluency until 80% in TeSR-E8TM medium and transferred to TeSR-E6TM Medium (StemCell Technologies), lacking Fibroblast Growth Factor 2 (FGF2), to promote spontaneous differentiation. To favor the differentiation into a neuroectoderm lineage, N2 supplement was added to the media 2 days after. On day 28, identified self-forming neural-like structures were isolated and transferred to 24 well plates in pro-neural medium (DMEM/F12, 1:1, L-Glutamine, 1% MEM nonessential amino acids, 2% B27 supplement) with 10 units/ml penicillin, and 10 mg/ml streptomycin. 7 days later, FGF2 was removed. At day 84, retinal organoids werre cultured in pro-neural medium with 2% B27 supplement without vitamin A. The media was changed every 2-3 days during all the differentiation.

### Mouse model

All animal experiments were realized in accordance with the NIH Guide for Care and Use of Laboratory Animals (National Academies Press, 2011). The protocols were approved by the Local Animal Ethics Committees and conducted in accordance with Directive 2010/63/EU of the European Parliament. The project was evaluated by the CEEA 05 (Ethical Committee in Animal Experimentation 05) and approved by the MESRI (“Ministère de l’Enseignement Supérieur, de la Recherche et de l’Innovation”, France). The approval number of the projects from the animal facility is: B-75-12-02 and C-75-12-02. In this study, two adRP mouse models were used. The first one exhibits the *RHO* p.P347S mutation and has been published by Li and colleagues. (Li et al. 1996) The second model used harbors the *RHO* p.P23H mutation and was published by Robichaux and colleagues. (Robichaux et al. 2022 Mar 11) In each case, the mutation is not located in the sgRNA on-target site to avoid any inherent editing variability between them.

### RNP lipofection on HEK293T cells

HEK293T cells were plated in 48 well-plates at 1E6 cells per well at a total volume of 250μL per well. 24 hours after plating, cells (2E6 cells per well, 70% confluency) were lipofected with SpCas9 RNP at a final concentration of 100nM and mixed with Lipofectamine 2000 (Invitrogen) according to the manufacturer’s instructions. To ensure reliability, each SpCas9 RNP complexed with one specific sgRNA was lipofected in three different wells (internal replicates) and the experiment was replicated three times (independent repeats).

Briefly, sgRNA and SpCas9 (Aldevron) were mixed at 1:1 molar ratio by pipetting and incubated at room temperature for 5 minutes. Then, RNP were mixed with Lipofectamine 2000 (Invitrogen) and Opti-MEM (ThermoFisher), vortexed for 10 seconds and incubated at room temperature for 10 minutes. Lipofection mix containing SpCas9 RNP was then added to corresponding wells for a total final volume of 275 µL. Lipofected cells were harvested 48 hours after lipofection, DNA was extracted using NucleoSpin Tissue Kit (Macherey-Nagel) according to the manufacturer’s protocol and conserved at -80°C for further experiments.

### « Live » imaging of the HEK293T lipofection

HEK293T cells were plated in 24 well plates at 1E5 cells per well at a total volume of 400 μL per well. The same protocol was used using a SpCas9-GFP. 25 μL of lipofection mix containing SpCas9-GFP RNP were added to corresponding wells after 24 hours for a final volume of 425 μL. For control wells (non-lipofected cells), 25 μL of Opti-MEM was added to the wells.

45 minutes post-lipofection, cells were observed under DMI6000b epifluorescent microscope Leica Microsystems) at 20X and images were captured every 15 minutes for 42 hours. The microscope used for live imaging was equipped with an incubation chamber to maintain the cells at 37°C and 5% CO₂ throughout the experiment. Image analysis was performed using ImageJ. Videos were edited using ImageJ and Cap Cut.

### RNP lipofection in retinal organoids in 96 well plate

We calculated to have the same amount of Cas9 RNP mixed with Lipofectamine 2000 for the same number of cells. *In vitro*, we plated 1E5 cells per well in a 48 well plate. 24h later, when we started the transfection, we had 2.16E5 cells per well. For the organoids, at D150, we counted 1.26E5 cells per organoid. As we transfected 100 nM of Cas9 RNP in HEK293T, to keep the same ratio of Cas9 RNP per cell, we used 60 nM of Cas9 RNP with 20% Lipofectamine 2000 per organoid. The same RNP/Lipo mix procedure than for HEK lipofection was applied.

### Naked Cas9 RNP treatment of retinal organoids using a hanging drop

sgRNA and SpCas9 (Aldevron) were mixed at 1:1 molar ratio by pipetting and incubated at room temperature for 5 minutes. Then, RNP were diluted in culture media. Each organoid was placed on the lid onto the cover of a 35mm petri dish, and excess culture media was removed. 6 µL of RNP/media was placed as a drop onto the organoids. The bottom of the petri dish was covered with culture media to ensure it stayed moist. The cover was turned upside down and the organoids were placed into the incubator overnight. The next morning, the organoids were transferred back to 24 ultra-low attachment well plates, with 500 µL of fresh culture media per organoid per well and were placed into the incubator.

### AAV design, production and serotypes used

The AAV construct encoding for SpCas9 was designed as published by Wu and colleagues (Wu et al., 2022). SpCas9 was driven by a sCMV promoter. The second AAV construct encoding for sgRNA was designed with sgRNA under the control of U6 promoter and a GFP reporter cassette under the control of CAG ubiquitous promoter. AAV plasmids were purchased from VectorBuilder.

AAV vectors were produced as previously described using the co-transfection method and purified by iodixanol gradient ultracentrifugation. (Gray et al. 2011) AAV vector stocks were titrated by quantitative polymerase chain reaction (qPCR) (Aurnhammer et al. 2012) using PowerSYBR Green (Thermo Fisher Scientific). Retinal organoids were transduced with AAV 2.7m8 variant. Mouse retinas were transduced with AAV 8.

### Retinal organoids transduction of AAV

The AAVs were mixed with culture media to achieve the dosage of 1E9 vg per organoid. Retinal organoids were transferred from their 24 well-plates into a 96 well-plate in a total volume media of 50 µL. The plate was placed into the incubator overnight. The following morning, 100 µL of culture media was added to each well and the organoids were placed into the incubator. 24 hours later, the organoids were transferred back to 24 ultra-low attachment well plates, with 500 µL of fresh culture media per organoid per well and were placed into the incubator. Retinal organoids were collected 2 weeks after transduction.

### Subretinal injections in mice

Mice were anesthetized by isoflurane inhalation (Isorane, Axience). Pupils were dilated and 1 µL subretinal injections were performed at P14 using a Hamilton syringe with a 33-gauge blunt needle (World Precision Instruments) under an operating microscope (Leica Microsystems). Ophthalmic ointment (Fradexam) was applied after surgery. Eyes with extensive subretinal hemorrhage were excluded from the analysis. Animals were euthanized by CO2 inhalation and cervical dislocation. For editing analysis, the whole neural retina was isolated 7 days after injection (for RNP injection) or 3 weeks after injection (for AAV injection) and stored at -80°c prior to DNA/RNA extraction.

### Eye fundus imaging

Mice were anesthetized by isoflurane inhalation (Isorane, Axience). Pupils were dilated using 0.5% tropicamide (Mydriaticum®, Thea) and 5% phenylephrine hydrochloride (Neosynephrine®, Europhta). During the experiment, ophthalmic lubricant (Lubrithal®, Dechra) was applied. Micron IV (Phoenix Micron) was used to generate both bright field and fluorescence eye fundus images.

### Samples preparation for gene editing quantification

For HEK293T cells, genomic DNA was extracted using the NucleoSpin® DNA tissue (Macherey-Nagel) following the manufacturer’s instructions.

For neural retina tissue or human retinal organoids, DNA and RNA were simultaneously extracted from the same sample using the Quick-DNA/RNA Microprep Plus Kit (Ozyme) according to the manufacturer’s instructions.

PrimeSTAR GXL DNA polymerase (Takara) was used for PCR amplification according to the manufacturer’s instructions with a total of 50 ng of genomic DNA for each sample. Primers for the amplified region of interest are listed in supplemental tables S2 and S3. PCR amplicons were then purified using NucleoSpin® PCR and gel kit (Macherey-Nagel) following the manufacturer’s instructions.

### Sanger sequencing and TIDE analysis on HEK293T

To detect CRISPR/Cas9-mediated gene editing on targeted loci for each sgRNA, primer couples were designed to amplify on-target sites according to Tracking of Indels by Decomposition (TIDE) website recommendations (http://shinyapps.datacurators.nl/tide/). Purified PCR amplicons were sent to Eurofins Genomics Europe for Sanger sequencing. Sequences were analyzed using TIDE algorithm to estimate the spectrum and frequency of indels from both sides of the theoretical cleavage site (sgRNA sequences were implemented in analysis). Results given by the algorithm were expressed as the total estimated editing percentage.

### Next-generation amplicon sequencing for organoids and mouse retinas

For mouse retina and retinal organoids experiments, purified PCR amplicons were sent to Next Generation Sequencing (NGS) platform at Massachusetts General Hospital DNA core facility (https://ccib.mgh.harvard.edu/). Interleaved Fastq files were then submitted to CRISPResso2 analysis. (Clement et al. 2019) Briefly, sgRNA 5’-3’ spacer and amplicon sequences were uploaded prior to analysis. Classical analysis parameters remained unchanged. Editing percentages from the first generated figure expressing modified reads were conserved as suggested by CRISPResso2 creators.

### Reverse Transcriptase-Quantitative PCR

Organoids and retina were collected respectively, 2 weeks or 3 weeks after transduction or injection. RNA was extracted using the Quick-DNA/RNA Microprep Plus Kit (Ozyme) according to the manufacturer’s instructions and submitted to TURBO DNAse (Invitrogen) treatment according to the manufacturer’s instructions. Reverse transcription (RT) was performed on the extracted DNA-digested RNA samples using Superscript Reverse Transcriptase IV (Invitrogen) following the manufacturer’s instructions with oligodT or random hexamers (for sgRNA) primers (Thermo Fisher Scientific). qPCR was performed using *Power*SYBR Green PCR Master Mix (Thermo Fisher Scientific) according to the manufacturer’s instructions. A total of 5ng of cDNA was used per well, in duplicate. Duplicates that had a Ct standard deviation above 0.5 were removed. For semi-quantitative analysis, Ct values were converted into relative expression scores using the formula 35 - Ct, where 35 was used as the threshold value to determine target expression. For all conditions, the mean value from untreated samples was subtracted to normalize expression levels. Negative values resulting from this normalization were set to 0. Undetermined Ct values from untransduced samples were set on Ct=35.Primers used are listed in Table S4.

### Cytotoxicity assay

CyQUANT™ LDH Cytotoxicity Assay Kit (Invitrogen) was used to analyze cytotoxicity, following the manufacturer’s instructions.

The supernatant medium of floating organoids was collected 7 days post-treatment and transferred to a 96-well flat-bottom plate and LDH release was evaluated. Absorbance was read using Spark microplate reader (TECAN) at 490 and 680 nm and the 680 nm absorbance value (background signal) was subtracted from the 490 nm absorbance value to determine LDH activity.

### DLS

Stunner (Unchained Labs) was used to measure the AAV empty vs full ratio and for dynamic light scattering (DLS) measurements. A 2 μL sample was loaded onto specialized 96-well plates, which include microfluidic channels and dual-cuvettes (Unchained Labs). For AAV quantification, the AAV Quant application was selected in the Lunatic & Stunner Client software and for Cas9 RNP quantification, the protein classic Q280 application was selected. Data were acquired using 4 acquisitions of 5 sec. Data analysis was performed using the Lunatic & Stunner Analysis Software.

### Statistical analysis

All statistical analyses were carried out using GraphPad PRISM version 7.0. P-values were determined by ordinary One-way ANOVA test, Dunnett’s multiple comparisons. ns: non-significant, * p<0.05, ** p<0.005, *** p<0.0005, **** p<0.0001.

### Immunostaining and imaging

1, 3 or 7 days post-infection, retinal organoids were fixed in a 4% formaldehyde solution for 1 hour. 1, 3 or 7 days post-injection, mouse eyes were enucleated and immediately fixed in a 4% formaldehyde solution for 1 hour.

To prepare cryosections, neural retinas were immersed in PBS-10% sucrose for 1 hour and then PBS-30% sucrose overnight at 4°C. They were embedded in OCT medium (CellPath) and frozen in liquid nitrogen. Retinal organoids were fixed in 4% PFA for 10 min at 4°C, cryoprotected in PBS with 30% sucrose overnight, embedded in PBS with 7.5% gelatin and 10% sucrose, and frozen in isopentane at -50°C. 10 μm-thick vertical sections were cut with a CM3050S cryostat (Leica Biosystems).

After 3 PBS washes, cryosections were incubated in a blocking buffer (For retinas: PBS, 1% bovine serum albumin, 0.1% Triton-X100, 0.1% Tween20. For retinal organoids: 0.2% gelatin and 0.1% Triton X-100) for 1 hour and then with primary antibodies diluted in blocking buffer and incubated overnight at 4°C. Primary antibodies used in this study are listed in Table S5. After three PBS washes of the sections, the secondary antibodies diluted in blocking buffer (Alexa Fluor 488, 594 or 647, Thermo Fisher Scientific) were added for 2 hours at room temperature, followed by three PBS washes and counterstained with DAPI. Cryosections were mounted in Vectashield mounting medium (Vector Laboratories) and visualized using a confocal microscope (Olympus). ImageJ software was used to process the images.

### Transgene copy number quantification

Transgene copy number in the Tg(p.P347S *hRHO*) mouse model was quantified using Droplet Digital PCR (ddPCR) following the protocol described by Bell et al. (Bell et al. 2018) The Tg(p.P347S *RHO*) mouse was generated by DNA microinjection in zygotes, resulting in the integration of an unknown number of transgene copies. Genomic DNA was extracted from tail tissue samples, and 10 ng of DNA was used for ddPCR analysis after digestion with the HindIII-HF restriction enzyme (New England Biolabs) according to the manufacturer’s protocol.

To quantify the transgene copy number, a multiplex ddPCR reaction was prepared using probes and primers designed for both the transgene and a reference locus, actb. The transgene-specific probe (h*RHO*-P347S-HEX) was labeled with HEX fluorophore, and the reference probe (actin-FAM) was labeled with FAM fluorophore. The ddPCR reaction mixture was prepared using Bio-Rad’s ddPCR Supermix for Probes (no dUTP) and droplets were generated using QX-100 Droplet generator. Amplification was run on C1000 Touch Thermal Cycler (Bio-Rad) with the following parameters: 95°C for 15min (2°C/Sec) then, 40 cycles at 94°C for 30s, 60°C for 1min, finished by 98°C°C for 10min. After amplification, droplets were analyzed using the Bio-Rad QX-100 Droplet Reader to calculate the transgene copy number relative to the reference gene actin. The probes and primers were designed using Primer3 (Table S6) and synthesized by Integrated DNA Technologies (IDT).

## Supporting information

Supplemental results

## Acknowledgements

We are grateful to C. Botto, Y. Rasool and V. Barrios Guttierez for preliminary in vivo and in vitro work, sample preparation and indel analysis, and to all the team S15 for their feedback on the manuscript and helpful discussions. JP was supported by grants from the Fondation pour la Recherche Médicale (FRM SPF201909009287) and the Foundation Fighting Blindness (PPA-0922-0840-INSERM). This work was supported by grants from AFM Trampolin Grant (Project 28641), ANR (ANR-23-CE18-0036-01 PolyCas9_RD), LABEX LIFESENSES [ANR-10-LABX-65], IHU FOReSIGHT (ANR-18-IAHU-01), INSERM, Sorbonne Université, Fondation Voir et Entendre and DIM Thérapie Génique and DIM C-BRAINS, funded by the Conseil Régional d’Ile-de-France

## Author Contributions

JP, HM, PO and DD designed experiments. ADC and MA produced Cas9 proteins according JPC supervision. HM designed sgRNAs and the AAVs. HM and PO performed transfection on cell lines. LV performed the imaging of the HEK293T cells. CR and EA produced the retinal organoids. CR, EA and PO performed the infections of the organoids. PO, EA and HM performed the Cyquant analysis. JP and HM performed sub-retinal injections in mice. JP, HM, PO and LV dissected mice eyes. JP, HM, PO, EA and LV extracted the DNA and prepared and analyzed NGS samples. ST, JP, HM, PO, LV, performed histology, immunostaining and imaging. JP performed and analyzed DLS recordings. HM performed the qPCR. JP and HM performed and analyzed the fundus images. OG brought the iPSCs and organoids culture protocol and provided the hCAR ab. JP, HM and DD wrote the manuscript. OG, JPC and DD provided scientific input and gave feedback on the manuscript.

## Data and materials availability

The data that support the findings of current study are available from the corresponding author upon reasonable request.

