## Supplemental results for "Retinal organoids mirror CRISPR/Cas9 gene editing efficiency observed in vivo"

**(See additional videos)**

**Supplemental Video S1: Timelapse imaging of non-transfected HEK293T cells.**
Non-transfected HEK293T cells were imaged over a 42-hour period. Cell growth and division are observed throughout the experiment. Live images were taken every 15min.

**Supplemental Video S2: Timelapse imaging of CRISPR/Cas9 transfection in HEK293T cells.**
Cas9-GFP RNP mixed with lipofectamine complexes (in green) interact with HEK293T cells over a 42-hour period after transfection. Capture of an RNP aggregate by a cell is highlighted in two zoomed-in sections. Live images were taken every 15min.

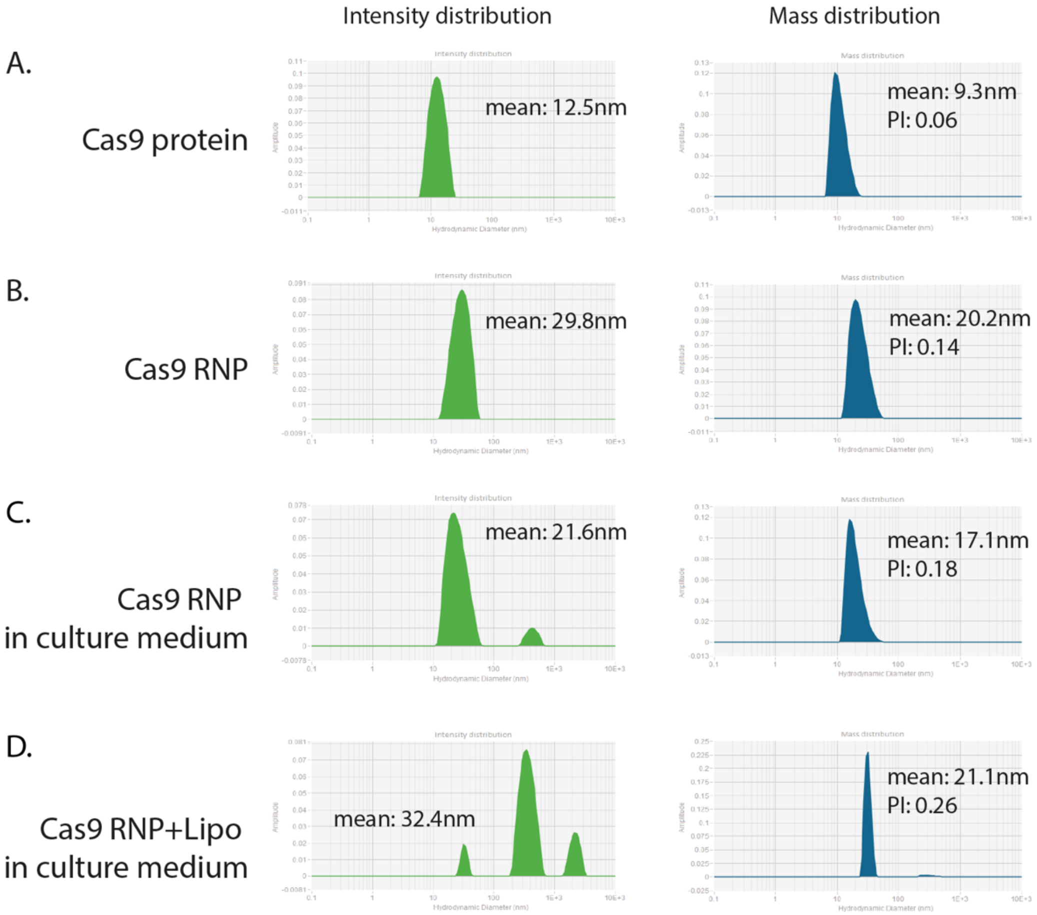


**Figure S1: Hydrodynamic size of Cas9 RNP and Cas9 RNP complexed with lipofectamine 2000 in culture medium.**

Size distribution by intensity and mass of Cas9 protein and its sgRNA: (A) of the Cas9 protein alone (B) of the Cas9 protein mixed with the sgRNA (C) of the Cas9 protein mixed with the sgRNA and incorporated in the culture medium (D) of the Cas9 protein mixed with the sgRNA and with Lipofectamine 2000 and incorporated in the culture medium.

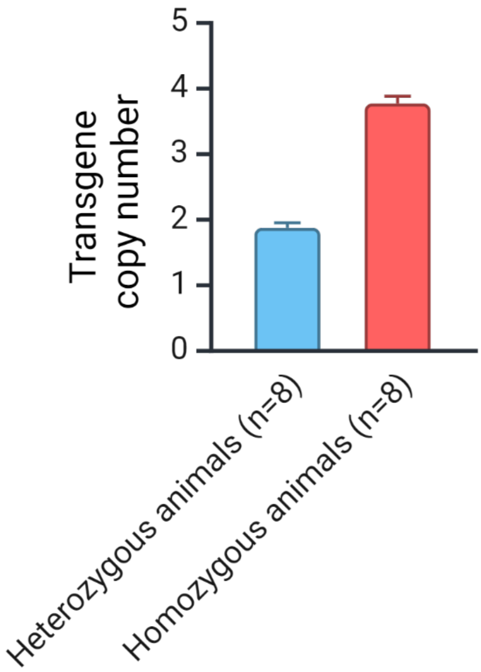


**Figure S2: Transgene copy number of RHO in the RHO.P347S mouse line quantified using Droplet Digital PCR (ddPCR)**

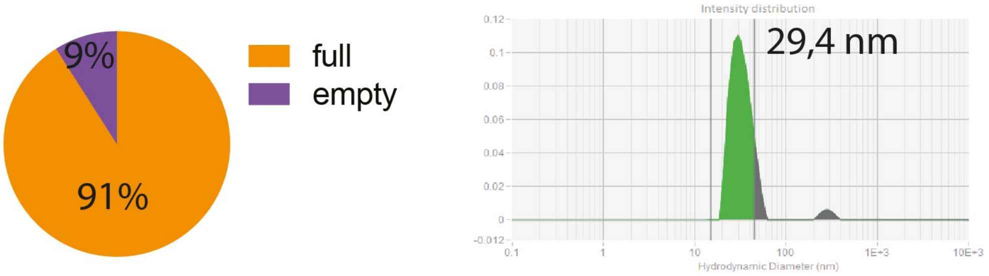


**Figure S3: DLS characterization of the SpCas9 AAV construct.** Left panel: pie chart showing the percentage of full (orange) vs empty (purple) AAV capsid. Right panel: hydrodynamic size distribution by intensity of AAV:SpCas9. Mean capsid diameter (nm).

| sgRNA name | Position on *RHO* | Spacer (5'-3') | PAM (5'-3') |
|---|---|---|---|
| **sgRNA 1** | Exon 1 | GATCAGCAGAAACATGTAGG | CGG |
| **sgRNA 2** | Exon 1 | TTGAGCAGGATGTAGTTGAG | AGG |
| **sgRNA 3** | Exon 1 | AGAGCGTGAGGAAGTTGATG | GGG |
| **sgRNA 4** | Exon 1 | AGCCACCTAGGACCATGAAG | AGG |
| **sgRNA 5** | Exon 1 | AGTACTGTGGGTACTCGAAG | GGG |
| **sgRNA 6** | Exon 2 | TGGCATGGTTCTCCCCGAAG | CGG |
| **sgRNA 7** | Exon 2 | CAGGACCACCAAGGACCACA | GGG |
| **sgRNA 8** | Exon 2 | TTGCAGGTGAAATTGCCCTG | TGG |
| **sgRNA 9** | Exon 3 | CTACTACACGCTCAAGCCGG | AGG |

**Table S1. sgRNA sequences used in this study**

| Primer name | Experiment | Sequence (5'-3') |
|---|---|---|
| Sanger-sgRNA1-Fw | TIDE analysis | GGG GTC AGA ACC CAG AGT CA |
| Sanger-sgRNA1-Rev | TIDE analysis | TGG CAA AGA AGC CCT CCA AA |
| Sanger-sgRNA2-Fw | TIDE analysis | GAA TGG CAC AGA AGG CCC TAA |
| Sanger-sgRNA2-Rev | TIDE analysis | CAT TGA CAG GAC AGG AGA AGG G |
| Sanger-sgRNA3-Fw | TIDE analysis | ATT CTT GGG TGG GAG CAG C |
| Sanger-sgRNA3-Rev | TIDE analysis | ACC CAC ACC CGG CTC ATA C |
| Sanger-sgRNA4-Fw | TIDE analysis | ATG GCA CAG AAG GCC CTA AC |
| Sanger-sgRNA4-Rev | TIDE analysis | TGA CAG GAC AGG AGA AGG GA |
| Sanger-sgRNA5-Fw | TIDE analysis | TCA GCC AGG AGC TTA GGA GG |
| Sanger-sgRNA5-Rev | TIDE analysis | TGC AGA GAG GTG TAG AGG GT |
| Sanger-sgRNA6-Fw | TIDE analysis | GGT TGC CTT CCT AGC TAC CC |
| Sanger-sgRNA6-Rev | TIDE analysis | TAC ACC CCT ACC CTG AGT GG |
| Sanger-sgRNA7/8-Fw | TIDE analysis | TTT CTT TGC CCA GCT CTC CT |
| Sanger-sgRNA7/8-Rev | TIDE analysis | TGA GTC CTG ACT GGA GGA CC |
| Sanger-sgRNA9-Fw | TIDE analysis | GCA TCT GCA TCC CCA TCT GA |
| Sanger-sgRNA9-Rev | TIDE analysis | GTC CAG ACC ATG GCT CCT C |

**Table S2. Primers used to amplify the targeted regions of each gene for Sanger sequencing and TIDE analysis**

| Primer name | Experiment | Sequence (5'-3') |
|---|---|---|
| NGS-sgRNA7-Fw | NGS-CRISPResso2 | TGC TCA GTG CCA TTA CCT GGA |
| NGS-sgRNA7-Rev | NGS-CRISPResso2 | GAG TGC ACC CTC CTT AGG C |
| NGS-sgTS2-Fw | NGS-CRISPResso2 | TGG GCT TCC CCA TCA ACT TC |
| NGS-sgTS2-Rev | NGS-CRISPResso2 | TGG CAA AGA AGC CCT CCA AA |
| NGS-sgRNA1-Fw | NGS-CRISPResso2 | CCC ACA GTA CTA CCTGGC TG |
| NGS-sgRNA1-Rev | NGS-CRISPResso2 | GGG CCC GAA GAC GAA GTA TC |

**Table S3. Primers used to amplify the targeted regions of each gene for NGS sequencing**

| | | |
|---|---|---|
| qPCR-sgRNA7-Fw | **sgRNA expression** | **CCA AGG ACC ACA GTT TTA GAG C** |
| qPCR-sgRNA7-Fw | sgRNA expression | CGG TGC CAC TTT TTC AAG TT |
| qPCR-SpCas9-Fw | Cas9 expression | AAC AGC CGC GAG AGA ATG AA |
| qPCR-SpCas9-Rev | Cas9 expression | CAC GGG GTG TTC TTT CAG GA |

**Table S4. Primers used for the RT-qPCR**

| Antigen | Dilution | Source |
|---|---|---|
| **DAPI** | 1/2000 | Thermo A31573) |
| **Recoverin** | 1/3000 | Millipore MAB 5585 |
| **hCAR** | 1/10.000 | Homemade. Already published in (*1*) |
| **RHO** | 1/500 | Merck MABN15 |
| **Peripherin** | 1/250 | Proteintech 18109-1-AP |
| **SpCas9** | 1/1000 | Genscript A01935-40 |

**Table S5. Primary antibodies used for immunostaining**

| **Probe hRHO-P347S-HEX** | **Transgene CNV** | **HEX/ CCT CTC AGA /ZEN/ CCC TCG CAG C** |
|---|---|---|
| **Probe actin-FAM** | Transgene CNV | FAM/ AGG CAG CCA /ZEN/ GGG CTG GC |
| **hRHO-CNV-Fw** | Transgene CNV | GGA GGA ATG AAT GGG AAG GG |
| **hRHO-CNV-Rev** | Transgene CNV | CCA GGG AGG GAA AAA CAA CT |
| **Actin-CNV-Fw** | Transgene CNV | TAT GAA GGC TTT GGT CTC CC |
| **Actin-CNV-Rev** | Transgene CNV | CAC AGA GCC ACA AGC TGT TT |

**Table S6. Primers used for the transgene copy number quantification**

### Refence

1. M. Garita-Hernandez, M. Lampič, A. Chaffiol, L. Guibbal, F. Routet, T. Santos-Ferreira, S. Gasparini, O. Borsch, G. Gagliardi, S. Reichman, S. Picaud, J.-A. Sahel, O. Goureau, M. Ader, D. Dalkara, J. Duebel, Restoration of visual function by transplantation of optogenetically engineered photoreceptors. *Nature Communications* **10** (2019).
